# Support for the deuterostome clade comes from systematic errors

**DOI:** 10.1101/2025.01.13.632777

**Authors:** Ana Serra Silva, Paschalis Natsidis, Laura Piovani, Paschalia Kapli, Maximilian J. Telford

**Affiliations:** Centre for Life’s Origins and Evolution, Department of Genetics, Evolution and Environment, University College London, Gower Street, London WC1E 6BT, UK; Science Group, Natural History Museum, London SW6 5BD, UK

## Abstract

There is a long-standing consensus that the animal phyla closest to our own phylum of Chordata are the Echinodermata and Hemichordata. These three phyla constitute the major clade of Deuterostomia. Recent analyses have questioned the support for the monophyly of Deuterostomia, however, showing that the branch leading to deuterostomes is very short and may be influenced by systematic error. Here we use a site-by-site approach to explore multiple sources of error. Under conditions that promote long-branch attraction (LBA) – especially branch-length heterogeneity and sites constrained in their amino acid composition – we find that deuterostome monophyly is strongly supported. When we make efforts to mitigate these sources of error, we cannot distinguish between monophyletic and paraphyletic Deuterostomia. Our findings have implications for the interpretation of putative deuterostome fossils, for the reconstruction of a bilaterian ancestor and, more generally, for how datasets for deep-time phylogenetic analyses are assembled and analyzed.

**Teaser:** The apparently close relationship between Chordata and Ambulacraria (echinoderms and hemichordates) is boosted by a long-branch attraction artefact.

## Introduction

Phylum-level relationships of the Metazoa have been hotly debated for over 150 years (*1*) but one constant feature, since it was proposed in 1908 (*2*) has been the clade of Deuterostomia. While there have been changes to the phyla included in Deuterostomia (the lophophorates (*3*) and chaetognaths (*4, 5*) are now recognized as protostomes), the long standing consensus is of a deuterostome clade containing the Chordata and Ambulacraria (Echinodermata and Hemichordata). The principal morphological synapomorphies that have been cited in support of Deuterostomia are deuterostomy (the blastopore becomes the anus), radial cleavage in the early embryo, enterocoelous mesoderm/coelom formation, and the presence of pharyngeal gill pores/slits (*6*). Alternative topologies that could relate Ambulacraria, Chordata and Protostomia (Fig. 1) have been named the Orthozoa tree (Protostomia plus Ambulacraria) and the Centroneuralia tree (Protostomia plus Chordata) (*7*).

**Fig. 1.**
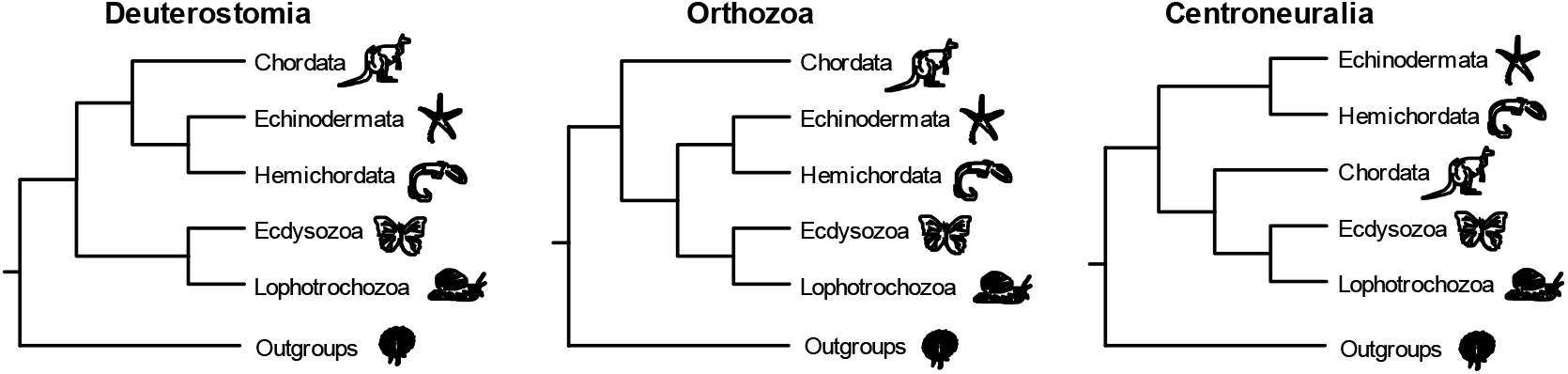
Topologies showing all possible relationships between deuterostome clades and Protostomia.

A modern conception of the animal tree of life, including the discovery that lophophorates and chaetognaths are protostomes, suggests that most of these embryological characters (deuterostomy, radial cleavage and enterocoely) are not deuterostome synapomorphies but plesiomorphies that were present in Urbilateria – the last common ancestor of bilaterians. From a morphological/embryological point of view, the deuterostomes can now be reliably defined only by their shared gill slits/pores (*7*) and possibly the presence of an endostyle (*8*). Recently it has been shown that the lack of robust support for deuterostome monophyly also extends to molecular data. In their study comparing molecular support for the two sister clades of Deuterostomia and Protostomia, Kapli et al. (*7*) found that out of a set of 31 genes originally believed to be unique to the deuterostomes (*9*) 24 could also be found in protostomes and/or non-bilaterian taxa, when using newer genome resources with better species sampling.

Searching the recent literature provides further evidence of weak support for Deuterostomia from molecular data; we have collated results from two decades of phylogenomic studies of metazoan relationships (*4, 10–24*) and found inconsistent support for the deuterostome branch. One trend we observe shows that studies that use site-homogeneous substitution models [e.g., (*10, 11, 15, 22, 24*)], typically, strongly support deuterostome monophyly, whereas studies that used site-heterogeneous models [e.g., (*4, 19, 22*)] showed weaker support for Deuterostomia or even supported paraphyletic Deuterostomia (Fig. 2). In some of the studies where site-homogeneous models were used, re-analyses of those datasets with site-heterogeneous models recovered paraphyletic Deuterostomia (*15–17, 22*). More generally, we notice that, even when using similar models, different studies conflict over their support for monophyletic deuterostomes suggesting the result is susceptible to changes in the particular set of loci or taxa that have been selected for study. Overall it seems that, despite being a widely accepted part of the animal phylogeny for over a century, Deuterostomia is a poorly supported clade.

**Fig. 2.**
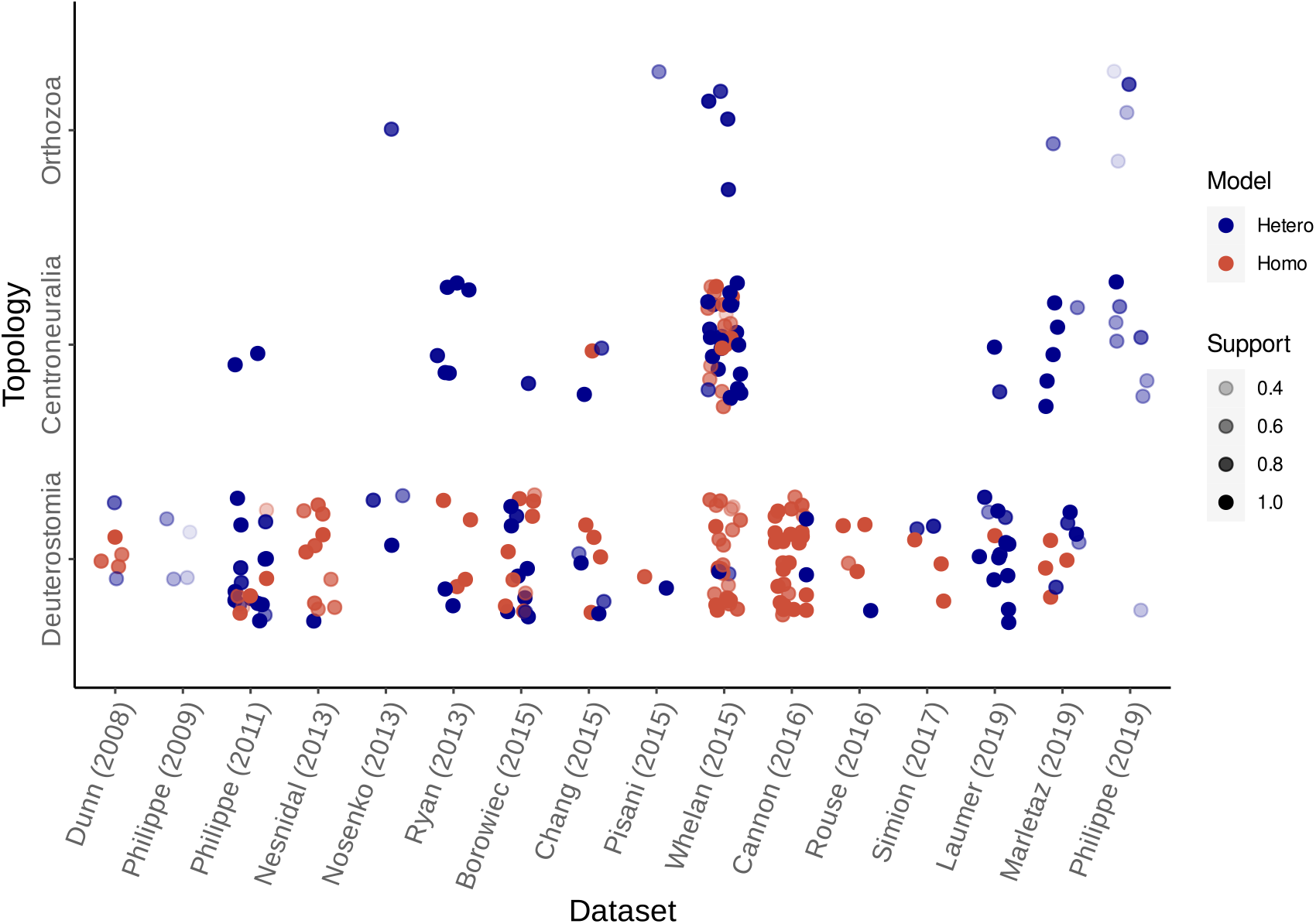
Normalized branch support for monophyletic Deuterostomia and its paraphyletic alternatives in the literature. Multiple studies on metazoan phylogenomics have recovered conflicting support for mono-or paraphyletic Deuterostomia. Support for deuterostome paraphyly is often observed in studies using site-heterogeneous substitution models (orange-red dots); whereas deuterostome monophyly is primarily supported by site-homogeneous models (blue dots). All branch support values (posterior probabilities, bootstrap and jackknife proportions) converted to proportions. Most studies used at least two branch support measures, commonly posterior probabilities and bootstrap proportions. The exceptions were (*14, 16, 20*), which used exclusively posterior probabilities, and (*22*), which used only bootstrap proportions.

Kapli et al. (*7*) re-analyzed five independently assembled phylogenomic datasets covering animal phyla (*4, 11, 14, 19, 21*) and showed that, in all of them, the internal branch leading to the deuterostomes was very short and, at best, weakly supported over the two alternative topologies. They also showed that, under conditions that might be expected to lead to a long-branch attraction (LBA) artefact in these datasets, support for deuterostomes was higher. The deuterostome branches of Chordata and Ambulacraria are not long-branched and the LBA artefact can be understood as an attraction between the long-branched Protostomia and the outgroup, leaving Chordata and Ambulacraria. clustered together to form a deuterostome clade. Consistent with this, Kapli et al. (*7*) simulated datasets according to the two possible trees on which the deuterostomes are paraphyletic – the Orthozoa (Ambulacraria+Protostomia) and the Centroneuralia (Chordata+Protostomia), see figure 1 – and found that conditions that emphasized LBA (unequal rates across branches and inadequate models) resulted in erroneous support for Deuterostomia.

Three important questions remain unanswered: first, how much does taxon selection contribute to the LBA artefact identified?; second, can we directly assess the effects of different possible sources of systematic error on the observed support for Deuterostomia using empirical data?; and third, do we still recover support for deuterostome monophyly in empirical data when the sources of identified error are minimized?

To address these questions, we have built a new dataset of 183 orthologous protein coding genes from 306 metazoan taxa. We have followed a stepwise approach to dissect the possible contributions of different sources of error to the support observed for Deuterostomia. Our large and taxonomically broad dataset allows us to use randomized taxon jackknifing to investigate the effects both of alternative taxon sampling and of unequal branch lengths. We have used site-by-site log-likelihoods to dissect the effects of different site-specific heterogeneities in the data. Using this method, we have examined the effects of site-rate heterogeneity, of site-compositional heterogeneity, and of using less well fitting substitution models. We have examined the impact of each potential source of error on support for deuterostome monophyly and investigated what level of support for Deuterostomia remained after we attempted to mitigate the sources of LBA we had identified by using optimal taxon sampling and the best fitting models.

## Results

### A. Building a large new empirical dataset to test causes of topological discordance

Our comparisons of past molecular analyses of metazoan phylogeny show inconsistent support for deuterostomes (Fig. 2) – the different results observed could be due to the use of different sets of loci, different taxa, different models or all three. Our new supermatrix of 183 loci from 306 species allows us to separate some of the effects of these parameters. We have used our large taxon sample to perform taxon jackknifing, subsampling species to explore how taxon sampling influences support for Deuterostomia for a single set of loci. Our site-by-site approach allowed us to study sampling effects across the alignment and to measure the support for different topologies in sites with different biases.

We collected the 306 metazoan proteomes and transcriptomes from various online databases (see dataProvenance.xlsx in DRYAD repository https://doi.org/10.5061/dryad.t76hdr89k) from species that represent the four major bilaterian branches of Ambulacraria, Chordata, Ecdysozoa and Lophotrochozoa, as well as a sample of non-bilaterian animals (see Table 1 and Materials and Methods). We used OrthoFinder (*25*) to identify 183 single-copy orthologs that were present in at least 70% of our 306 species (see Materials and Methods). We used trimAl (*26*) with option ‘-gappyout’ selected to remove unreliably aligned positions from each MAFFT (*27*) alignment and concatenated all 183 loci to make a supermatrix with 71,635 aligned amino acid positions and 69% average site occupancy. We built a maximum likelihood (ML) phylogenetic tree for all 306 species using IQ-TREE2 (*28*) under the LG+F+G4 model.

**Table 1.** Classes and sampling proportions for jackknifed alignments. Table showing the total number of non-deuterostome species sampled per class (fast-or slow-evolving) and per jackknife replicate. The first column shows the classes of taxa categorized by evolutionary rates(e.g., Lophotrochozoa Fast), with the following columns showing the total number of taxa sampled per class, the phyla in each class and the number of taxa sampled in each taxon jackknifing strategy. Long-branched jackknives included 12 deuterostomes and 12 ambulacrarians, for a total of 60 species per replicate. Short-branched replicates include an additional 13 deuterostomes and 13 ambulacrarians.

| Species classes | Number of species sampled | Phyla in species class | Number of taxa sampled in long-branch jackknives | Number of taxa sampled in short-branch jackknives |
| --- | --- | --- | --- | --- |
| Lophotrochozoa Fast | 20 | Platyhelminthes, Gastrotricha, Rotifera, Micrognathozoa | 12 | 0 |
| Lophotrochozoa Slow | 61 | Mollusca, Annelida, Brachiopoda, Nemertea, Bryozoa, Chaetognatha, Cyclophora | 0 | 13 |
| Ecdysozoa Fast | 38 | Nematoda | 12 | 0 |
| Ecdysozoa Slow | 41 | Arthropoda, Priapulida,<br>Tardigrada | 0 | 13 |
| Non-Bilateria<br>Fast | 13 | Porifera, Ctenophora | 8 | 0 |
| Non-Bilateria<br>Slow | 16 | Cnidaria | 4 | 8 |

The results of Kapli et al. (*7*) suggested that support for Deuterostomia could be inflated by an LBA artefact grouping Protostomia and long-branched species in the outgroup. To identify taxa in these two groups with higher or lower rates of evolution, we measured the average branch length per phylum using a custom python script (pynt.py) and classified any non-bilaterian, chordate and ambulacrarian phylum with an average clade branch length greater than 0.70 substitutions per site as fast-evolving. For phyla within the Lophotrochozoa and Ecdysozoa, we used higher branch length thresholds of 0.85 and 1.00, respectively.

Using the sets of protostomes and outgroups classified in this way as short-or long-branched, we generated two sets of taxon jackknifed alignments: 100 of our random taxon samples included both slow- and fast-evolving protostomes and outgroups; 100 used only slow-evolving protostomes and outgroups (Fig.3A). For each replicate, we randomly selected a fixed number of species (see table 1) from each of the four major branches of Bilateria (Ecdysozoa, Lophotrochozoa, Chordata and Ambulacraria) plus the outgroups. Each jackknife replicate contained 60 taxa in total and included all 71,635 alignment positions of the original alignment.

We first wanted to examine the effects of the long branches leading to protostomes and outgroups. We compared support for Deuterostomia across taxon jackknife replicates that included long-branched protostomes and outgroups (long-branched jackknives) with replicates excluding these (short-branched jackknives). We analyzed 100 long-branched and 100 short-branched jackknives using the LG+F+G4 model, which assumes homogeneous amino acid frequencies across the alignment (“site-homogeneous” hereon). Under LG+F+G4, all 100 of the long-branched jackknives supported Deuterostomia. Almost all short-branched replicates also support Deuterostomia, the exception is a single replicate that supported the Orthozoa topology. This very small difference might nevertheless hint at some degree of support for Deuterostomia deriving from branch length heterogeneities (Fig.3B).

We were interested in exploring two different potential origins of support for the three topologies (Deuterostomia, Orthozoa and Centroneuralia). First, we used the typical approach of comparing each topology’s log-likelihood score. We found (as discussed above) that all but one jackknife replicates supported deuterostome monophyly (table S1). Our second approach was to count the number of sites unequivocally supporting a single topology of interest over the other two. We did this by scaling the computed site log-likelihoods for each of the three topologies (such that they ranged from 0 to 1), and classifying any site with a scaled log-likelihood ≥ 0.66667 as supporting a single topology. We compared the numbers of such sites supporting each topology in both long- and short-branched jackknife replicates.

For long-branched jackknives, the Deuterostomia topology is always supported by many more sites (∼2.3x) than the two alternative topologies (Fig. 4A and table S1). For short-branched jackknives, however, the average number of sites preferring Deuterostomia roughly halves, while the average number of sites supporting the two paraphyletic Deuterostomia topologies change considerably less (average 23% increase in sites preferring Orthozoa and 4% decrease in sites preferring Centroneuralia; Fig. 4B). In 50 of the short-branched alignments there are more sites supporting Deuterostomia, in 49 a majority of sites support Orthozoa and in a single alignment there are more sites supporting Centroneuralia.

It is important to emphasize here that the strong effect of species sampling was only detected because we employed the taxon jackknifing approach. If we had used a single taxon sample (as is typical for phylogenomic studies), depending on the species represented we might have found a dataset with either a majority of sites preferring Deuterostomia or a majority supporting Orthozoa. This highlights both the fragility of the phylogenetic signal in these challenging relationships and the inherent weakness of typical phylogenomic methods that rely on a single instance of taxon sampling.

Our analyses show that many of the sites that strongly support Deuterostomia over the alternatives (i.e., scaled lnL ≥ 0.66667) do so only when long-branched protostomes and outgroups are included. Some of the support for deuterostome monophyly is therefore the result of unequal rates of evolution across branches, i.e. a long-branch artefact. This result supports Kapli et al.’s (*7*) observation that branch-length heterogeneity amplifies support for Deuterostomia, confirming the importance of careful taxon sampling to minimize LBA. We next wanted to test whether this branch-rate heterogeneity might be compounded by the effects of unaccounted for site-rate and site-compositional heterogeneities.

### B. Site homogeneous models support deuterostome monophyly across datasets

Many sites in an alignment are limited in the amino acids that can be tolerated rather than being free to sample the full amino acid residue space (*29, 30*). The more constrained a site is the more the frequency of amino acid substitutions at the site is underestimated, with the consequence that the frequency of convergence on long-branches is underestimated. The compositional constraint at a site can be represented as the effective number of amino acids at that site (k_eff_) and, when combined with unequal branch-lengths, the most constrained sites (lowest k_eff_) are expected to promote LBA (*30*).

We first assessed the effects of site compositional heterogeneity on the support for the three alternative topologies by comparing the results from site-homogeneous models (discussed in previous section) to equivalent results using site-heterogeneous mixture models. Site-heterogeneous mixture models are less prone to underestimating the frequency of substitutions, have been repeatedly shown to be better-fitting than site-homogeneous models in such deep phylogenies and to be less prone to LBA [e.g., (*7, 31*)].

Our site-by-site analyses showed that, for long-branched jackknives, when we use the site-heterogeneous EDM model, the average number of sites supporting Deuterostomia decreased by 23% and Orthozoa by 6.5%, while those supporting Centroneuralia increased by 6% (Figs. 3A,C and table S1). Deuterostomia was nevertheless supported by more sites than the alternatives in all but one of the 100 replicates, which supported Centroneuralia.

**Fig. 3.**
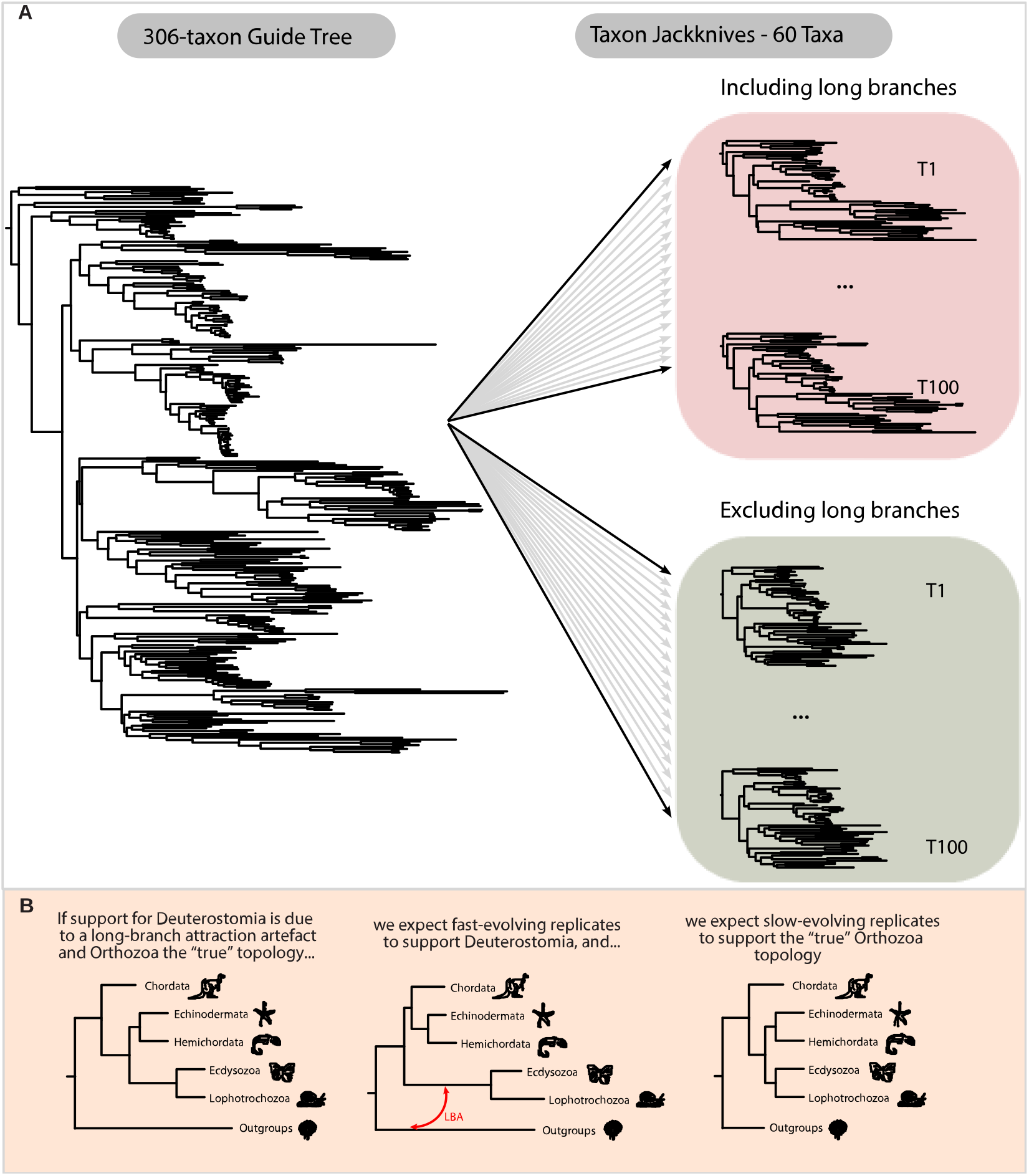
Summary of taxon jackknife protocol and possible effects of taxon-specific LBA on topology support. (**A**) Schematic representation of the randomized taxon jackknifing approach. (**B**) If the ‘true’ topology is one of the paraphyletic Deuterostomia alternatives; support for deuterostome monophyly results from a long-branch attraction artefact; but excluding long branch taxa should give the correct tree.

When we use the site-heterogeneous EDM model, for the short-branched jackknives in just over half of the replicates we found that more sites supported Orthozoa than Deuterostomia (Fig. 3B,D and table S1). In all jackknives far fewer sites supported Centroneuralia compared to the two other topologies. Compared to the long-branched replicates, the average number of sites supporting Deuterostomia decreased by 6%, those supporting Orthozoa decreased by 4%, and there was a negligible average increase of 0.3% sites supporting Centroneuralia.

Under the site heterogeneous empirical distribution mixture model [EDM+F+G4, (*29*)], the number of sites that do not prefer any of the three topologies increased noticeably in both short-and long-branched replicates compared to the analyses under LG+F+G4 (Fig. 4).

**Fig. 4.**
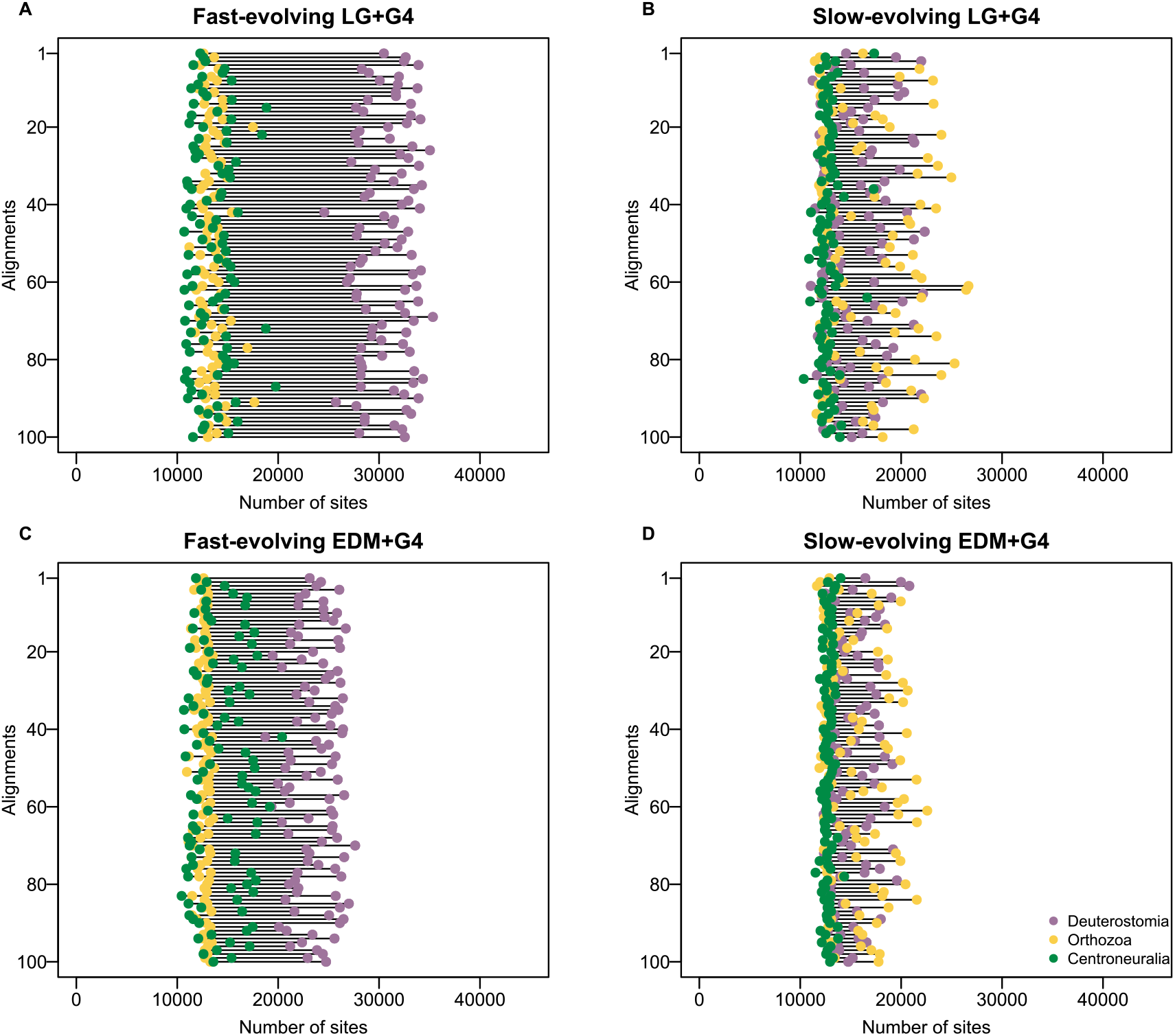
Monophyletic Deuterostomia is supported by fewer alignment sites under conditions avoiding long-branch attraction. Counts of sites preferring each of the three topologies using site-likelihood scores. (**A**) All 100 fast-evolving taxon jackknife replicates analyzed under the site-homogeneous LG+G4 model show a clear majority of sites (c.30,000) supporting deuterostome monophyly. (**B**) For 100 slow-evolving replicates, support for monophyletic Deuterostomia is dataset-dependent with approximately 50% of all jackknife replicates having a majority of sites supporting Deuterostomia and the other half having a majority of sites supporting the Orthozoa topology. (**C**) Under the finite-category site-heterogeneous EDM model, a smaller majority of sites (c.25,000) in the fast-evolving jackknives support Deuterostomia. (**D**) Under the EDM model slow-evolving replicates retain the 50/50 split between support for Deuterostomia or Orthozoa. Purple diamonds correspond to the Deuterostomia topology, yellow diamonds to Orthozoa and green diamonds to Centroneuralia.

Overall, using a site-heterogeneous model reduces the average number of sites supporting Deuterostomia more than the numbers of sites supporting the two paraphyletic topologies; this is emphatically true of the long-branched replicates. The implication is that a considerable proportion of the sites supporting Deuterostomia under site-homogeneous models are LBA-prone sites.

### C. Highly constrained amino acid sites support Deuterostomia

To explore further the effects of site-compositional heterogeneity on support for monophyletic Deuterostomia, we estimated the effective number of amino acids (k_eff_) for each site in our alignments (100 long-branched and 100 short-branched jackknife replicates). We assigned each site to a k_eff_ bin (from 1 to 20), following the protocol in Szánthó et al. (*30*). For each site in each k_eff_ bin we then plotted the difference in site log-likelihood observed between each possible pair of topologies (Deuterostomia vs Orthozoa; Deuterostomia vs Centroneuralia; Orthozoa vs Centroneuralia).

For all taxon jackknives and models tested, we found that support for Deuterostomia (over either of the other topologies) was greatest in the most highly constrained sites (lowest k_eff_) and that the degree of support for Deuterostomia decreased with increasing k_eff_ (Figure 5 A-B,D-E). For sites that have higher numbers of effective amino acids (k_eff_ > 15) the three alternative topologies (Fig. 5 A-B,D-E) had roughly equal support. The preference for Deuterostomia at sites with low k_eff_ values is reduced both when long-branched taxa are excluded (short-branched jackknives) and, as expected, when a site-heterogeneous model is used. When we compare support for the two paraphyletic topologies, under both site homogeneous LG+F+G4 and site heterogeneous EDM+F+G4 and under all taxon sampling combinations, we find that Centroneuralia is very slightly more strongly supported than Orthozoa, with the difference slightly more marked in sites with lower k_eff_ (Fig. 5 C,F).

**Fig. 5.**
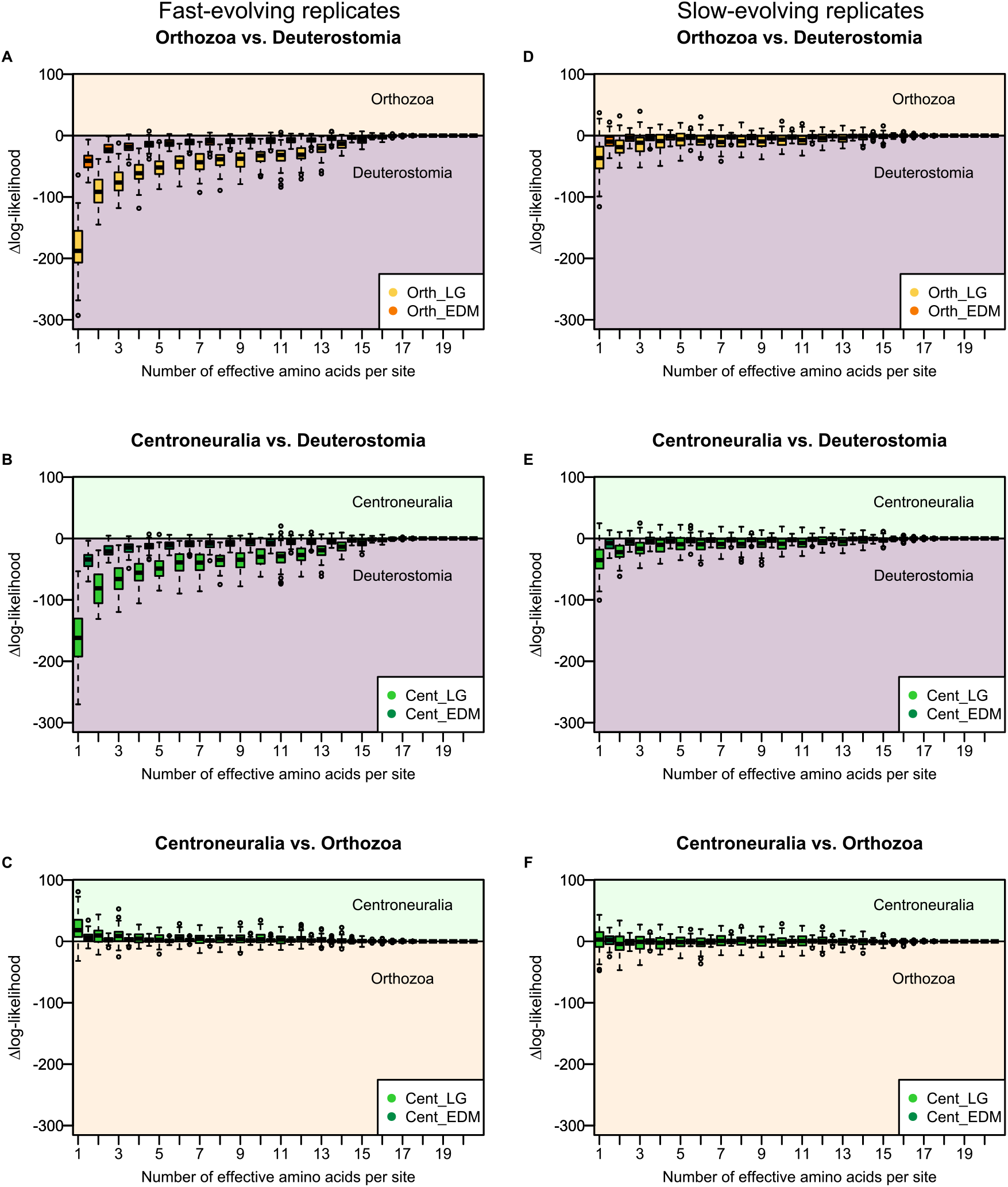
Highly constrained sites support deuterostome monophyly. The site-specific Δlog-likelihood between every pair of topologies, binned by their effective amino acid number (k_eff_) and under the site-homogeneous LG+F+G4 and site-heterogeneous EDM+F+G4 models. For the fast-evolving jackknife replicates, Deuterostomia is supported (**A**) over Orthozoa and (**B**) over Centroneuralia, especially at sites with low k_eff_ and under the site-homogeneous model. Comparing the two paraphyletic topologies (**C**) shows limited support for Centroneuralia over Orthozoa. For the slow-evolving jackknife replicates, Deuterostomia is still, if less strongly, supported (**D**) over Orthozoa and (**E**) over Centroneuralia. The comparison between the paraphyletic topologies (**F**) still supports Centroneuralia at low k_eff_ values, but for most bins the support is either very low or does not distinguish between the two topologies. For all plots, boxes in the light purple area support Deuterostomia, in the salmon Orthozoa, and in the green area support Centroneuralia; Yellow boxes (**A, D**) represent the distribution of Δlog-likelihoods between Orthozoa and Deuterostomia under the LG+F+G4 model, and orange boxes the distribution under the EDM+F+G4 model. Bright green boxes represent the distribution of Δlog-likelihoods between Centroneuralia and Deuterostomia (**B** and **E**), or between Centroneuralia and Orthozoa (**C** and **F**), under the LG+F+G4; and dark green boxes show the same for analyzes under EDM+F+G4. The black bars correspond to the median Δlog-likelihoods.

We have shown that the most highly constrained sites (which are expected to contribute most to LBA errors) are indeed those that most strongly support Deuterostomia. We note that using a model (EDM) that is designed to accommodate site-compositional heterogeneity reduces but does not eliminate all support for Deuterostomia. The EDM model has a finite set of categories of sites (here 128 categories) and there remains the possibility that it cannot fully accommodate all effects of site-compositional heterogeneity.

Rapidly evolving sites are also expected to be prone to LBA artefacts, even if they have repeatedly been shown to retain phylogenetic signal [eg., (*32*)]. We repeated all analyses without the site-rate variation parameter (LG+F and EDM+F) and found that, overall, support for monophyletic Deuterostomia increased under site-rate homogeneous models (see Supplementary Materials). More specifically, the number of sites supporting Deuterostomia increased in both fast- and slow-evolving jackknife replicates when we did not use a gamma correction (Figs.4 and S1-3).

### D. Infinite sites model cannot distinguish between mono- and paraphyletic Deuterostomia

Finite-category mixture models like EDM were developed as less computationally expensive approximations of the infinite categories CAT model [e.g.,(*29, 30*)]. When applied to real data, however, these mixture models are typically worse-fit models than CAT and can be less effective at mitigating LBA artefacts [e.g., (*7*)].

To test how well the EDM model mitigates LBA artefacts in our data, we simulated 40 alignments under fixed (and therefore known) Deuterostomia and Orthozoa topologies using the infinite sites CAT-Poisson+G4 model in PhyloBayes (*33*). Simulation parameters were initially estimated using the CAT-Poisson+G4 model and a under one of two fixed topologies (either Deuterostomia or Orthozoa). We used the posterior distribution of parameters estimated in this way to produce our simulated alignments. We simulated 40 alignments and analyzed these under the LG+F+G4 and EDM+F+G4 models as before (for detailed methods and results see Supplementary Materials). We found that the site-heterogeneous model (EDM+F+G4) always supports the ‘true’ topology (Deuterostomia or Orthozoa), whereas the less well fitting site-homogeneous model (LG+F+G4) nearly always supports deuterostome monophyly even when the underlying tree is Orthozoa. If a paraphyly-supporting short-branched empirical alignment is used to simulate data under the Orthozoa tree, then LG supports Orthozoa.

While the site-heterogeneous EDM+F+G4 model always supports the ‘true’ topology, suggesting that it sufficiently mitigates LBA, we wanted to compare the fit of the EDM+F+G4 model and the CAT model to real data and then to ask whether the slight overall preference for monophyletic Deuterostomia we observe with empirical data using EDM is also seen using the infinite-categories CAT model.

Inference under the CAT model is computationally expensive, especially for large datasets, and is prone to convergence and mixing problems (*34*), making it impractical to use it to repeat our topology tests across all 200 jackknifed replicates. We opted, instead, to compare the support for each of the three topologies under increasingly complex models (site-homogeneous LG, finite categories site-heterogeneous EDM and infinite sites site-heterogeneous CAT). We used ‘leave-one-out cross-validation’ analyses to make these comparisons. These model/topology comparisons used the empirical jackknife dataset that yielded the shortest tree under a Deuterostomia topology and the EDM+F+G4 model, for each jackknifing strategy. These alignments were chosen because they minimized branch-length heterogeneity arising from taxon sampling and so were expected to be the least likely to be affected by LBA artefacts. The Poisson exchangeability was chosen for the CAT model to match the exchangeability used to generate the EDM model(s) [following (*29*)]. To contextualize the results, we also used the same approach to compare support for topologies with monophyletic versus (highly improbable) paraphyletic Protostomia [Deuterostomia+Ecdyzosoa, P2 in Kapli et al. (*7*)].

We found that the CAT model was the best-fit model regardless of jackknife replicate (short-or long-branched) and, as expected, the topologies with monophyletic Protostomia were all better-fitting than paraphyletic Protostomia under all models and jackknife strategies. For the long-branched replicate, monophyletic Deuterostomia was always the preferred topology, regardless of model. However, the level of support was inversely proportional to model complexity.

For the short-branched alignment, monophyletic Deuterostomia was preferred over the two paraphyletic trees under both LG+G4 and EDM+G4 (table 2). The best fitting CAT-Poisson+G4 model, however, could not distinguish between the Deuterostomia, Orthozoa and Centroneuralia topologies. These results indicate that, under conditions that constitute our best attempt to minimize LBA (minimal branch length heterogeneities and site compositional heterogeneity dealt with using the CAT model), none of the three topologies is preferred over the other two.

**Table 2.**
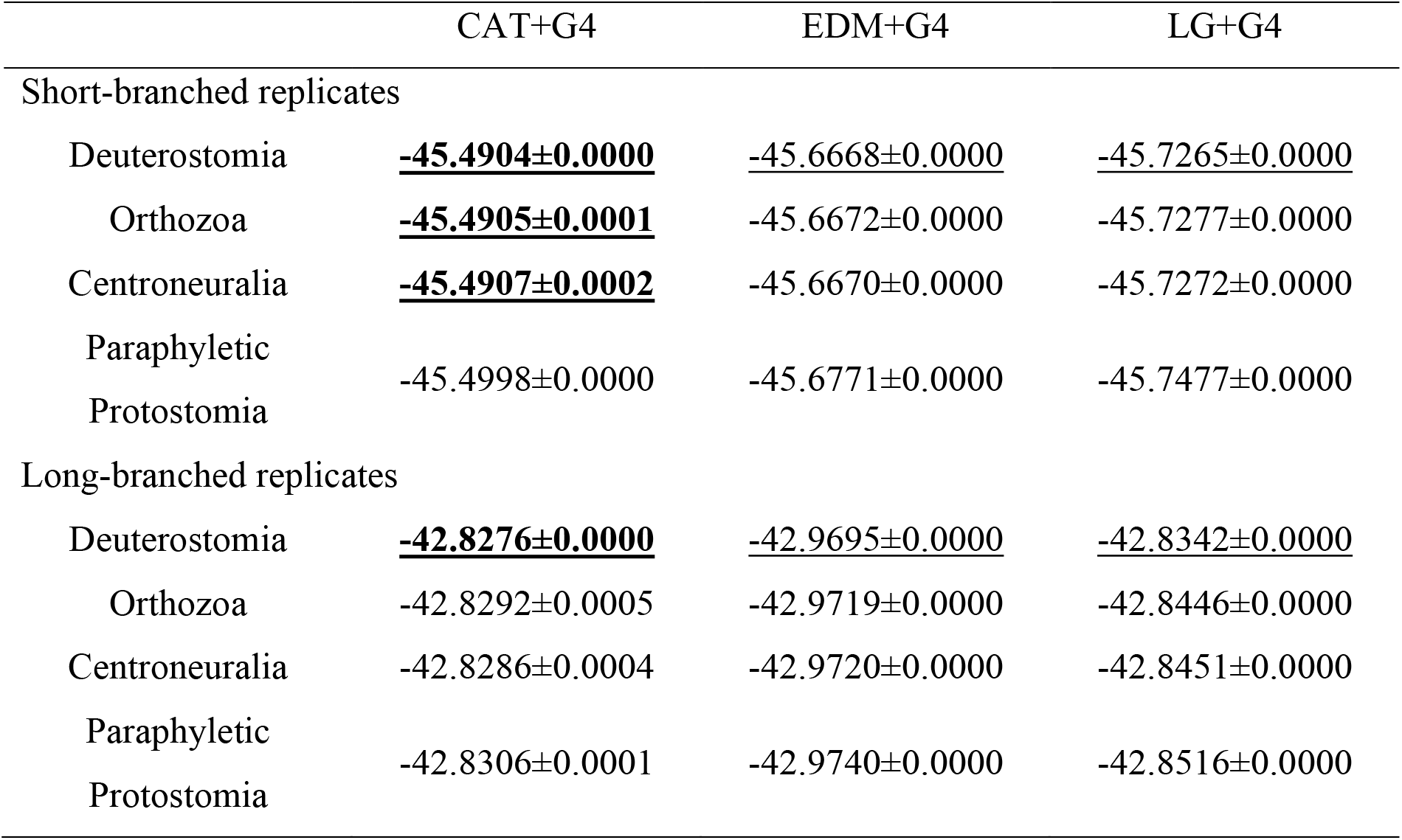
When sources of systematic error are minimized leave-one-out cross-validations cannot distinguish between mono- and paraphyletic Deuterostomia. Cross-validation analyses on the shortest short-branched alignment still support monophyletic Deuterostomia under a site-homogeneous (LG) and finite-category site-heterogeneous (EDM) models. However, under the best-fitting infinite-category site-heterogeneous CAT-Poisson model, we cannot distinguish between mono- and paraphyletic Deuterostomia. For the shortest of the long-branched alignments, Deuterostomia is always the preferred topology. The Deuterostomia, Orthozoa and Centroneuralia topologies all display monophyletic Protostomia. Underlined values correspond to the best-fit topology under each model and bold values to the overall best-fit topology per alignment.

## Discussion

The deuterostome clade, since it was proposed in 1908 (*2*), has been one of the few consistently accepted branches of the metazoan phylogeny. It has a special significance from an anthropocentric view as, if real, it reveals the closest relatives of our own clade of Chordata and so is important in efforts to reconstruct the earliest events in chordate and vertebrate evolution (*35*). The relationships of the deuterostome clades and the reconstruction of their common ancestor is also essential for the interpretation of certain Cambrian fossils with supposedly deuterostomian features [in particular those fossils interpreted as having pharyngeal slits, which are clearly homologous between Chordata and Ambulacraria (*36*)].

Our deeply engraved perception of animals as either “deuterostomes” or “protostomes” may explain why the monophyly of deuterostomes has not been more critically examined, despite accumulating evidence from phylogenetic studies over the past 15 years [e.g., (*4, 12, 14, 17, 19, 22, 24*)]. Our past and present work (*7*) highlights this long-overlooked but significant issue, demonstrating that deuterostome relationships should be as contentious as other debated branches in animal phylogeny, such as the placement of Ctenophora [e.g., (*13, 20, 22–24, 37*)], Xenacoelomorpha [e.g., (*11, 19, 21, 37*)] and Placozoa [e.g., (*38–42*)].

The work of Kapli et al. (*7*) provided tangible but still circumstantial evidence that support for Deuterostomia may derive from a LBA artefact linking Protostomia to the outgroup(s). The work described here provides thorough empirical and simulation-based evidence in support of the idea that Deuterostomia is supported by an LBA artefact. Furthermore, we have attempted to uncover the true relationships among the deuterostome clades (Chordata and Ambulacraria) and protostomes by meticulously addressing and mitigating identified sources of error.

We have shown that a combination of unequal branch-lengths and unaccommodated site-compositional heterogeneity strongly bias towards recovering monophyletic deuterostomes even though the true support for this clade is very weak. When we deal with the LBA artefact by removing long-branched protostomes and outgroups, our taxon jackknifing approach, as well as showing much weaker support for Deuterostomia, also revealed the great sensitivity of the support for Deuterostomia to the taxon sampling. This would not have been apparent from a typical single-sample study, nor from cross-dataset comparisons. The observed variability in these results reinforces the idea that, once sources of error are minimized, there is little and questionable support left for monophyletic Deuterostomia.

Our simulation experiments, though restricted in the potential sources of error modeled, showed that the finite-category mixture model we used (EDM) was able to reconstruct the correct tree even in the context of cross site-compositional heterogeneity. While this might seem to suggest that the overall support for Deuterostomia in the EDM analyzes confirms the null hypothesis that they are monophyletic, the inability of the better fitting CAT model to distinguish between topologies leaves the split between Chordata, Ambulacraria and Protostomia closer to a polytomy.

Our efforts to identify and minimize sources of systematic error on the earliest bilaterian splits though extensive, were not exhaustive. Sources of confusion whose impact we have yet to explore are those of branch-compositional heterogeneity and of loci with distinct evolutionary histories. In recent years there has, however, been a proliferation of models developed to model branch-composition heterogeneity [e.g., NDCH2 (*43*); GHOST (*44*)], site-branch-composition heterogeneity [GFmix (*45*)] and multi-tree histories [MAST (*46*)], which if combined with our jackknifing and site-by-site focused approach might shed further light on the deuterostome monophyly problem.

Evidence for non-tree like processes, however indirect, would lend further support to a (near) polytomy at the base of Bilateria and would therefore directly impact our interpretation of the last common ancestor of these lineages. As the time between successive cladogenesis events must have been very short (leaving little room for the evolution of novel characters) and the branches leading to crown Protostomia, Chordata and Ambulacraria are longer (allowing clade specific character loss) Urbilateria can be best reconstructed as having possessed any features in common to any pair of these three clades. This would mean, for example, that the pharyngeal slits common to Chordata and Ambulacraria are plausible characteristics of Urbilateria with obvious implications for our interpretation of Cambrian fossils with this character.

At this stage in the development of phylogenomic methods, it remains challenging to determine whether the apparent polytomy results from model limitations in capturing the complex heterogeneities of substitution processes, unresolved gene-tree/species-tree conflicts due to hidden paralogy [e.g., (*19, 47, 48*)], or potential non-tree-like processes, such as hybridization and incomplete lineage sorting, arising from a near-simultaneous diversification event [e.g., (*47, 49, 50*)]. Newer models—including those that address branch-compositional heterogeneity, site-branch heterogeneity, and multi-tree histories—show promise, but our understanding of their effectiveness for deciphering ancient divergences remains limited. Until these models are thoroughly validated through simulation-based assessments, their application to deep phylogenies should be approached with caution. In the meantime, the relationships among the three bilaterian clades of interest may be best represented as unresolved, reflecting the rapid and complex evolutionary processes likely underlying this foundational divergence.

## Materials and Methods

### Experimental Design

We used a combination of empirical and simulated datasets to test the effects of unequal rates of evolution, site-rate and site-compositional heterogeneity on the support for deuterostome monophyly. First, we used taxon jackknifing to generate subsampled alignments to test the effect of long-branched taxa on the support for monophyletic Deuterostomia. We then compared the support for deuterostome monophyly under site-homogeneous (known to exacerbate LBA artefacts) and better-fitting finite-category and infinite-sites site-heterogeneous models.

To test whether support for Deuterostomia is being amplified by or due solely to LBA, we then simulated data under a selection of the jackknifed alignments, a site-heterogenous model with infinite site categories and two guide trees–monophyletic and paraphyletic Deuterostomia.

### Data collection

We collected 306 metazoan genomic datasets from various databases [e.g., (*51–54*)], prioritizing broad taxonomic sampling and species with high genome coverage (accessions available from DRYAD repository https://doi.org/10.5061/dryad.t76hdr89k). The 306 animal proteomes sufficiently represented the four major bilaterian branches–79 Ecdysozoa, 82 Lophotrochozoa, 63 Chordata and 53 Ambulacraria–and 29 non-Bilateria (Table 1). The majority of selected proteomes contained genes predicted based on reference genomes, except for Ambulacraria data, where genomic sampling is relatively poor. We increased ambulacrarian sampling with high-quality transcriptomic data.

For the RNA data downloaded from NCBI SRA database, we assembled the transcriptomes using trinity v.2.13.2 (*55*) and predicted protein sequences using TransDecoder v.5.5.0 (https://github.com/TransDecoder/TransDecoder), both under default parameters. We assessed the completeness of the 306 proteomes with the single-copy orthologues in the BUSCO metazoan gene library (*56*), using a 55% cut-off. Lastly, we ran sample sequences belonging to the same NCBI BioProjects though CroCo v.1.1 (*57*) to screen for contaminants and removed flagged sequences.

### Orthology inference and supermatrix construction

We inferred gene orthology with OrthoFinder v.2.5.1 (*25*), under default settings and stopping the pipeline after the orthogroup inference step (“-og” flag). We used the OGFilter.py script (https://github.com/pnatsi/OGFilter.py) to select orthogroups present in at least 214 species (∼70% taxon coverage) and with at most seven paralogues/isoforms per species. This filtering yielded 183 multi-paralogue orthogroups.

The filtered orthologous gene sequences were aligned with MAFFT v.7.455 (*27*) under default settings. For each orthogroup, we inferred a maximum likelihood phylogenetic tree in IQ-TREE2 (*28*) with ModelFinder (*58*) enabled for the WAG, JTT and LG amino acid substitution models. We ran the resulting multi-paralogue orthogroup trees through the ParaFilter script (https://github.com/pnatsi/ParaFilter) to generate single-copy orthogroup trees/alignments suitable for phylogenetic inference. The ParaFilter script checks the relative position of all paralogues of a target species in the orthogroup tree and, if the paralogues form a clade (in-paralogues), keeps the shortest branch as the species’s representative sequence or, if paralogues are paraphyletic in the tree (out-paralogues), discards all sequences of the target species.

We re-aligned the resulting single-copy orthogroups with MAFFT and ran them through trimAl v.1.4.rev15 (*26*) with “-gappyout” selected to remove gaps. The trimmed alignments were concatenated into a supermatrix with 306 taxa and 71,635 amino acid residue sites, which was run through IQ-TREE under the site-homogeneous LG+F+G4 model.

### Stratified random alignment subsampling

Beyond their taxonomic ranks, the 306 sampled species can be further classified into slow- and fast-evolving, based on their average clade branch-lengths measured from the maximum likelihood tree inferred from the concatenated matrix (available from DRYAD repository https://doi.org/10.5061/dryad.t76hdr89k). The average branch lengths were computed with the pynt.py script (https://github.com/MaxTelford/XenoCtenoSims/blob/master/pynt.py). Non-protostome phyla with average branch lengths greater than 0.70 were classed as fast-evolving. The thresholds for Ecdysozoa and Lophotrochozoa were 0.85 and 1.00, respectively. Monotypic phyla were randomly assigned to the fast-or slow-evolving categories. The sampled species were divided into ten categories (five major taxonomic groups and two evolutionary rates), as shown in Table 1.

To investigate the effect of taxon sampling on the support for monophyletic Deuterostomia, we generated subsamples of species from the 306-species matrix. Each jackknifed alignment contained 60 species, with pre-defined proportions of the 10 taxon/rate categories mentioned above (Table 1). Two different category proportion settings were applied: one containing only slow-evolving Protostomia and outgroups (‘short-branch’ datasets); and one containing both slow- and fast-evolving Protostomia and outgroups (‘long-branch’ datasets). This jackknifing strategy was designed to test for a possible (positive) correlation between inclusion of long-branched protostome and outgroup taxa and support for monophyletic Deuterostomia. Fast-evolving chordates and xenambulacrarians were omitted from both subsampling strategies to restrict the problem to deuterostome monophyly. In total, 200 alignments of 60 species each were drawn from the 306-species supermatrix–100 ‘short-branched’ and 100 ‘long-branched’.

### Testing for the effects of branch lengths and site-rate heterogeneity on supported topology

We assessed the phylogenetic support for the Deuterostomia, Orthozoa and Centroneuralia topologies across all 200 taxon jackknifed alignments by comparing site-specific log-likelihood scores under fixed topology (‘-te’) searches in IQ-TREE2 and four substitution models: a site-homogeneous model with or without discrete gamma categories and a site-heterogenous substitution model with or without gamma.

The guide trees (available from DRYAD repository https://doi.org/10.5061/dryad.t76hdr89k) were obtained by pruning three 306-taxon trees, each representing a topology of interest (Deuterostomia, Orthozoa or Centroneuralia) to the taxa present in each subsampled alignment. The 306-taxon trees were constructed by subtree prune-regrafting of the inferred ML 306-taxon tree to make the inferred phylum-level relationships congruent with the most up-to-date phylogenomic studies (*4, 14, 19*).

For the site-homogeneous analyses we selected the LG+F model, with the ‘+G’ runs set to the default 4-category discrete gamma distribution. For the site-heterogeneous analyses we used Schrempf et al.’s (*29*) empirical distribution mixture (EDM) model, which approximates Lartillot’s (*59*) CAT model in a maximum likelihood framework. The EDM profile was obtained from the PhyloBayes-MPI v.1.9 (*33*) computed site profiles (‘readpb_mpi -ss’) for the 306-taxon supermatrix and converted to an IQ-TREE-compatible input with the EDCluster tool (*29*) set to 128 clusters (‘-k 128’) and the log centred log-ratio transformation (‘-t LCLR’). In IQ-TREE, the EDM model was set as ‘-m Poisson+EDM0128LCLR+F+G4’.

All fixed-topology IQ-TREE2 searches were run with the ‘-wslr --rates’ selected, which outputs a file with site-specific log-likelihoods and a file with site-specific rate categories. IQ-TREE assigns each site into a rate category based on its evolutionary rate using an empirical Bayesian approach (*60*); slowest sites are assigned to category 1, and fastest sites to category 4. To count the number of sites supporting each topology of interest, we scaled the inferred site-specific likelihood scores with the custom likelihood_transform.py pyhton script (https://github.com/MaxTelford/MonoDeutData/tree/master/scripts/likelihood_transform.py) and set the threshold for single-topology supported to a scaled lnL ≥ 0. 0.66667.

We used the effective number of amino acids per site (k_eff_) measure to explore how support for the three alternative topologies (Deuterostomia, Orthozoa and Centroneuralia) changes across sites with different levels of amino acid residue constraint. The k_eff_ of each site was computed based on the same site profile used to generate the EDM model exchangeability rates, using the scripts provided in Szánthó et al. (*30*). We performed three pairwise support comparisons of the IQ-TREE runs: Deuterostomia *vs*. Orthozoa, Deuterostomia *vs*. Centroneuralia and Orthozoa *vs*. Centroneuralia (Fig. 5)

### Leave-one-out cross-validation analyses

Following the recommendation in Lartillot (*61*) and the PhyloBayes manual, we used PhyloBayes-MPI to run multiple leave-one-out cross-validation tests on the shortest empirical jackknifed alignment with and without long-branched taxa. We tested four topologies–Deuterostomia, Orthozoa, Centroneuralia and a paraphyletic Protostomia– and three models (LG, EDM and CAT-Poisson) with and without gamma (+G4), for a total of 24 cross-validation tests per alignment.

## Acknowledgments

We thank members of the Telford and Yang labs (UCL) for discussions, Jeff Thompson for sharing a large number of Ambulacraria genomes with us prior to their publication and Daniel Leite for his transcriptome assembly scripts.

## Funding

Leverhulme Trust Research Grant RPG-2021-433 (MJT/ASS)

BBSRC grant BB/R016240 (MJT/PK)

European Union’s Horizon 2020 research and innovation program under the Marie

Sklodowska-Curie grant agreement no 764840 IGNITE (MJT/PN)

Marie Skłodowska-Curie (grant agreement number 766053 EvoCELL (MJT/LP)

## Author contributions

Conceptualization: MJT, PK

Methodology: ASS, PN, PK

Investigation: ASS, PN

Data curation: PN, LP

Visualization: ASS, PN

Supervision: MJT

Writing—original draft: ASS, PN, LP, PK, MJT

Writing—review & editing: ASS, PN, LP, PK, MJT

Funding acquisition: MJT

## Competing interests

All authors declare they have no competing interests.

## Data and materials availability

All data are available in the main text, the supplementary materials or the DRYAD repository https://doi.org/10.5061/dryad.t76hdr89k.

## Notes

https://doi.org/10.5061/dryad.t76hdr89k

### Competing Interest Statement

The authors have declared no competing interest.

