## Supplementary materials for "Support for the deuterostome clade comes from systematic errors"

Serra Silva *et al.*

**This PDF file includes:**

Supplementary Text
Figs. S1 to S6
Table S1

**Other Supplementary Materials for this manuscript include the following:**
Data S1

### Supplementary Text

Section 1 – Effect of discrete gamma rate categories on support for monophyletic Deuterostomia

The effects of unaccounted for branch-length and site-compositional heterogeneity on support for deuterostome monophyly were discussed extensively in the maim text; however, we also explored the effects of unaccounted for site-rate heterogeneity on the support for monophyletic Deuterostomia. Kapli et al. (*7*) have previously shown that models not accounting for site-rate heterogeneity are generally worse fitting than models that include this parameter and that inference under the worse fitting models affects branch-length estimation. Work on other recalcitrant branches of the Tree of Life has shown that rapidly evolving sites are more prone to long-branch attraction (LBA) artefacts than more slowly evolving sites but that these fast sites still retain phylogenetic signal (*31*).

*Site-rate heterogeneity-aware models vs. rate-naïve models*

Following the taxon-jackknifing strategy described in the main text's 'Materials and Methods' section, we found that, just as in the analyses modelling site-composition heterogeneity (Fig. 4), the exclusion of fast-evolving taxa considerably decreases the number of sites supporting monophyletic Deuterostomia (Fig. S1), as does the use of the empirical distribution site-heterogenous model [EDM, (*29*)]. It is worth noting, however, that while accounting for rate-heterogeneity with the site-homogeneous LG model leads to a decrease in the number of sites unequivocally supporting a single topology, the EDM model shows an opposing trend, where modelling rate-heterogeneity leads to an increase in single-topology supporting sites.

The per category (pseudo-category in the case of analyses not modelling rate-heterogeneity) trends are similar for the jackknifing replicates including fast-evolving taxa, with the number of sites supporting monophyletic Deuterostomia generally higher under the site-homogeneous LG+F model, except for pseudo-category 4, where the number of sites supporting Deuterostomia is nearly identical to those supporting the Orthozoa and Centroneuralia paraphyletic hypotheses. In pseudo-category 3 there is also an increase in the number of sites supporting the paraphyletic deuterostome topologies compared to rate category 3 (Fig. S2). As for the EDM model, more sites support Deuterostomia in all pseudo-rate categories compared to the analyses under the EDM+F+G4 model.

In the jackknife replicates excluding fast-evolving taxa there is no clear trend in changes in support for deuterostome monophyly between the true rate categories (in the +G4 models) and the pseudo-categories. Compared to the LG+F+G4 model, pseudo-categories 1 and 2 show higher numbers of sites supporting both Deuterostomia and Orthozoa and, much as in the fast-evolving replicates, pseudo-category 4 shows nearly identical numbers of sites supporting all three topologies of interest (Fig. S2). Pseudo-category 3, however, does show an increase in the number of sites supporting Orthozoa but a considerable decrease in sites supporting Deuterostomia, with several alignments where Deuterostomia is the topology with the fewest supporting sites. For the slow-evolving replicates, pseudo-rate categories 2-4 under EDM+F show little change from the rate categories under EDM+F+G4, but pseudo-category 1 shows a decrease in sites supporting Deuterostomia and Orthozoa.

Plotting topological support by (pseudo-)rate category, we see that for the jackknife replicates including fast-evolving taxa categories 1 and 4 favour Deuterostomia, while 2 and 3 tend to support the paraphyletic alternative, under both site-homogeneous and -heterogeneous models with little change in median delta log-likelihood between the models with or without Gamma (Fig. S3). Between paraphyletic topologies, under LG (pseudo-)categories 1 and 2 cannot distinguish between topologies, 3 favor Orthozoa and the fastest sites prefer Centroneuralia. Under the EDM model, (pseudo-)categories 2 and 3 cannot distinguish between alternatives, whereas category one supports Centroneuralia and 4 Orthozoa.

For short-branched replicates, there is little difference between the models with or without the Gamma parameter. Under the site-homogeneous model (pseudo)-category 1 cannot distinguish between the three tested topologies, category 3 supports the paraphyletic topology and 4 supports monophyletic Deuterostomia. Category 2 supports Deuterostomia over Orthozoa, but Centroneuralia over Deuterostomia. When comparing between support for the paraphyletic alternatives (pseudo-)categories 1 and 2 cannot distinguish between them, category 3 supports Orthozoa and 4 Centroneuralia. Under the EDM model (pseudo-)category 1 supports Orthozoa over Deuterostomia, but 2-4 support deuterostome monophyly. Compared to Centroneuralia, categories 1 and 3 support Centroneuralia and 2 and 4 support Deuterostomia. Between paraphyletic topologies, (pseudo-)categories 1 and 2 support Orthozoa, while 3 and 4 support Centroneuralia. Thus, unlike with the site-constrained effective amino acid [$k_{eff}$, (*28*, *29*)] analyses, where highly constrained sites always prefer monophyletic deuterostomes, we cannot link any (pseudo-)rate category to overwhelming support for Deuterostomia.

While it is clear that accounting for site-rate heterogeneity has an effect on the number of sites supporting monophyletic Deuterostomia, the changes are not unidirectional. When looking at differences in log-likelihood, however (pseudo-)category 4, the fastest-evolving, sites always show support for monophyletic Deuterostomia (Figs. S3).

*Testing reliability of slowest and fastest rate category sites with simulated data*

To gain a better understanding of how reliable sites in gamma distribution category 1 vs. sites in category 4 are, we generated simulated datasets using PhyloBayes v.1.9 (*32*) and the CAT-Poisson+G4 model. For each category, we simulated 100 alignments under the Deuterostomia and the Orthozoa topologies, with 30,000 sites each– (2 site rate categories) x (2 topologies) x (100 replicates). Each alignment was then used to infer a tree under the site-homogeneous LG+F+G4 model in IQ-TREE (*27*). For each of the four configurations (2 topologies x 2 categories), the resulting 100 trees were summarized with a greedy consensus tree.

The results (Fig. S4) show that the slow category 1 sites correctly recover the correct topology 100% of the time when the data were simulated across either the Deuterostomia or the Orthozoa topology. For alignments simulated under the fast-evolving category 4 sites, when the data were simulated across the Deuterostomia tree, it was correctly recovered 98% of the time. When the data were simulated across the Orthozoa tree, however, a Deuterostomia tree was incorrectly recovered 72% of the time. The fast-evolving sites were generally much more prone to error (and indeed errors were observed across both Deuterostomia and Orthozoa trees. We note that with real data, the fast sites are much more likely than slow sites to be adversely affected by alignment errors; with the simulated data we have eliminated this source of error though of course alignment errors will still have had an effect on the parameters estimated.

Section 2 – Finite site profile mixture models mitigate but do not negate LBA

*Untangling the effects of branch lengths and guide topology with simulated data*

To test how much choice of guide tree influenced the support for monophyletic Deuterostomia, we generated multiple simulated alignments with PhyloBayes under the CAT-Poisson model and parameters that match those measured from the empirical jackknifed alignments (site composition, per site amino acid diversity, etc.). Two alignments were chosen from the jackknifed alignments with and without long-branched taxa, the alignment with the highest and lowest relative support for monophyletic Deuterostomia based on the sums of site log-likelihoods. For each of the four alignments, PhyloBayes was run under a fixed topology (Deuterostomia and Orthozoa) until convergence was reached and 5 alignments were generated from the posterior distribution ('readpb_mpi -ppred'). After site profile and EDM exchangeabilities computation, the simulated alignments were re-run through the same IQ-TREE2 and $k_{eff}$ analyses as the empirical 60-taxon jackknifed alignments, see main text.

To see how well the EDM model mitigates LBA artefacts we simulated data under fixed (and therefore known) Deuterostomia and Orthozoa topologies using the infinite sites CAT-Poisson+G4 model in PhyloBayes. We modelled the simulations on parameters measured from four empirical jackknife replicates. The first two were the long-branched alignments that showed the strongest and weakest support for Deuterostomia. The second pair were the short-branched alignment showing the strongest support for Deuterostomia and the single alignment that supported deuterostome paraphyly (under LG+F+G4).

For each of the four empirical datasets, we estimated the model parameters (site composition, per-site amino acid diversity, etc.) according to fixed Deuterostomia and Orthozoa topologies. For each topology/alignment pair, we used the estimated parameters to generate five simulated alignments that spanned the posterior predictive distribution, and which had the same number of amino acid sites and taxa as the original jackknife alignments. For each simulated dataset, we measured the difference in support for Deuterostomia or Orthozoa across $k_{eff}$ bins when using the LG+F+G4 and EDM+F+G4 models.

For datasets simulated from each of the two selected long-branched alignments, the LG+F+G4 model always supported monophyletic Deuterostomia, regardless of the topology (Deuterostomia or Orthozoa) the data were simulated under. The EDM+F+G4 model, however, always supported the 'true' topology the data were simulated under (Figs.S5A, B and S6A,B).

For data simulated from the short-branched alignments, EDM+F+G4 still always supported the 'true' topology the data were simulated under (Fig.S5C, D and S6C, D. The site-homogeneous LG+F+G4 model supported monophyletic Deuterostomia, except for data simulated according to the paraphyly-supporting empirical alignment where LG+F+G4 supports the (correct) Orthozoa topology (Fig. S6D). In summary, site homogeneous models tend to incorrectly prefer the

Deuterostomia tree over the (correct) Orthozoa tree especially in the context of branch-length heterogeneity.

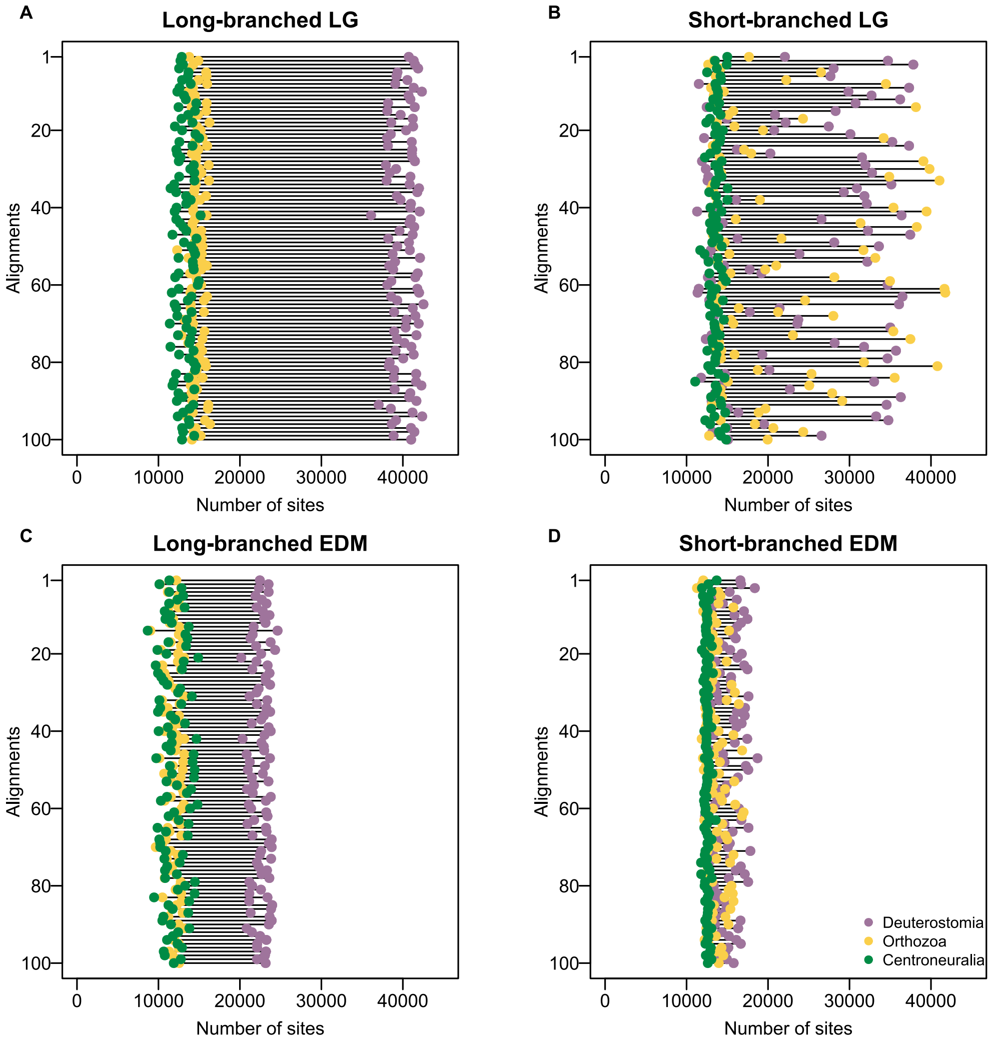


**Fig. S1.**

From site-likelihood scores, (**A**) all fast-evolving taxon jackknife replicates analyzed under the site-homogeneous LG model show a majority of sites (c.40,000) supporting deuterostome monophyly. (**B**) The same analyses, for the slow-evolving replicates show that support for monophyletic Deuterostomia is dataset-dependent with approximately 50% of all jackknifes supporting Deuterostomia and the other half supporting the Orthozoa topology. (**C**) Under the finite-category site-heterogeneous EDM model, the fast-evolving jackknifes still predominantly support Deuterostomia, but at lower numbers of sites (c.25,000). (**D**) Under the EDM model slow-evolving replicates retain the 50/50 split between support for Deuterostomia or Orthozoa but there is a drastic reduction in the number of sites supporting a single topology. Purple diamonds correspond to the Deuterostomia topology, yellow to Orthozoa and green diamonds to Centroneuralia.

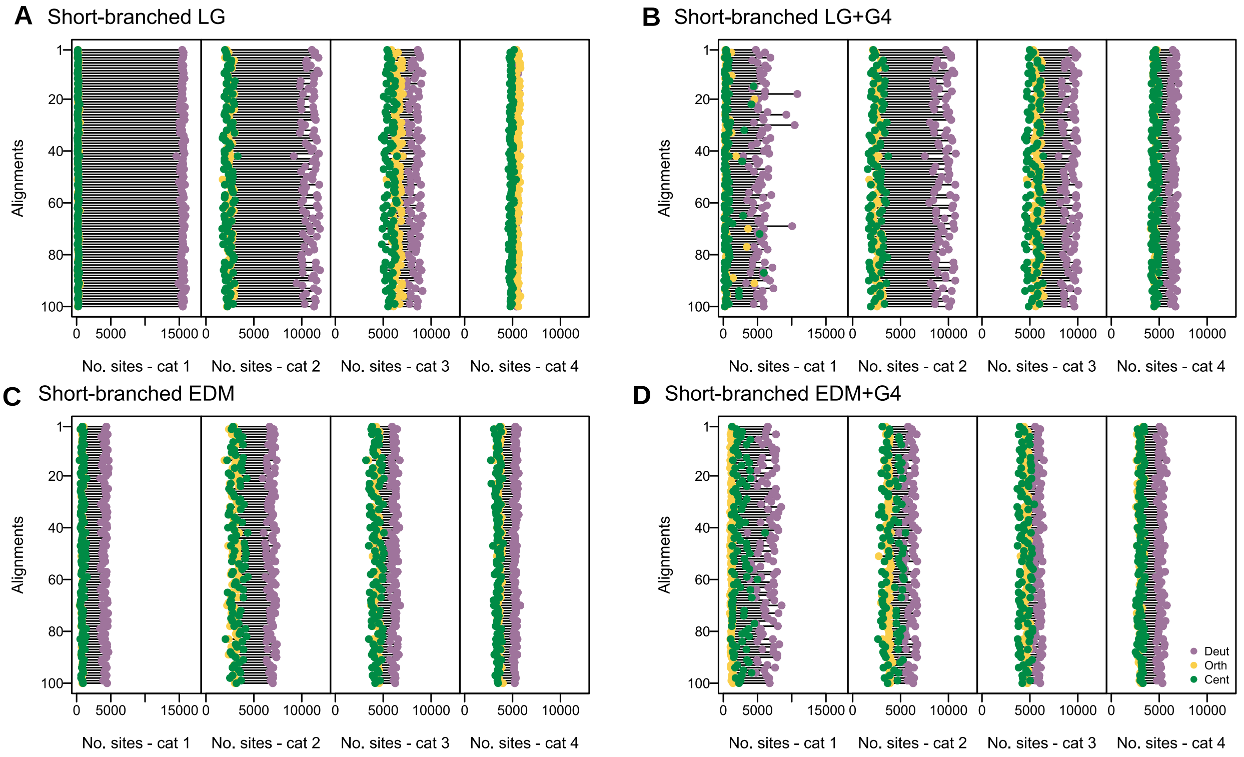


Fig. S2.

From site-likelihood scores and assignment of sites in the rate-naïve analyses to rate categories based on the '+G4' runs, (**A**) we see that for rate (pseudo-)categories 1 and 2, the slowest evolving sites, support for Deuterostomia is higher in the rate-naïve analyses under the site-homogeneous LG model. However, in the LG+G4 analyses (**B**) support for Deuterostomia is higher in the fastest evolving sites, (pseudo-)categories 3 and 4. Under the site-heterogeneous EDM model, there is a clearer separation of the amount of sites supporting monophyletic Deuterostomia versus its paraphyletic alternatives under the rate-naïve analyses (**C**). Under EDM+G4(**D**), monophyletic Deuterostomia is still favored in all rate categories, but less strongly in categories 2 and 3. Purple diamonds correspond to the Deuterostomia topology, yellow to Orthozoa and green diamonds to Centroneuralia.

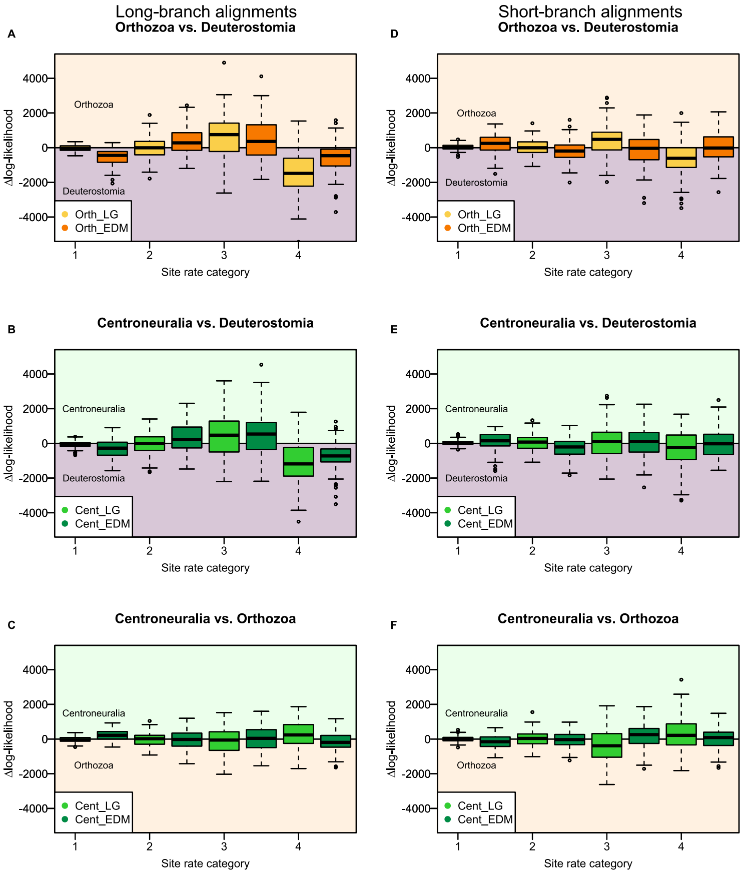


**Fig. S3.**

We show the site-specific Δlog-likelihood between every pair of alternative deuterostome topologies, binned by their rate categories and under the site-homogeneous LG+G4 and site-heterogeneous EDM+G4 models. For the fast-evolving jackknife replicates, Deuterostomia is, generally, supported over (**A**) Orthozoa and (**B**) Centroneuralia by sites in the fastest (4) and slowest (1) categories but sites in categories 2 and 3 tend to support paraphyletic Deuterostomia. Comparing the two paraphyletic topologies (**C**) shows limited support for Centroneuralia over Orthozoa by sites in category 1 under the site-heterogeneous EDM+G4 model and category 4 under LG+G4; However, category 4 sites support Orthozoa for analyses under EDM+G4. For the slow-evolving jackknife replicates, there is no clear pattern between support for mono- or paraphyletic Deuterostomia across models and rate categories – comparison to (**D**) Orthozoa and (**E**) Centroneuralia. The comparison between the paraphyletic topologies (**F**) shows a similar pattern to the long-branched replicates. For all plots, boxes in the light purple area support Deuterostomia, in the salmon Orthozoa, and if bars are in the green area they support Centroneuralia; Yellow boxes (**A**, **D**) represent the distribution of Δlog-likelihoods between Orthozoa and Deuterostomia under the LG+G4 model, and orange boxes the distribution under the EDM+G4 model. Bright green boxes represent the distribution of Δlog-likelihoods between Centroneuralia and Deuterostomia (**B** and **E**), or between Centroneuralia and Orthozoa (**C** and **F**), under the LG+G4; and dark green boxes show the same for analyzes under EDM+G4. The black bars correspond to the median Δlog-likelihoods.

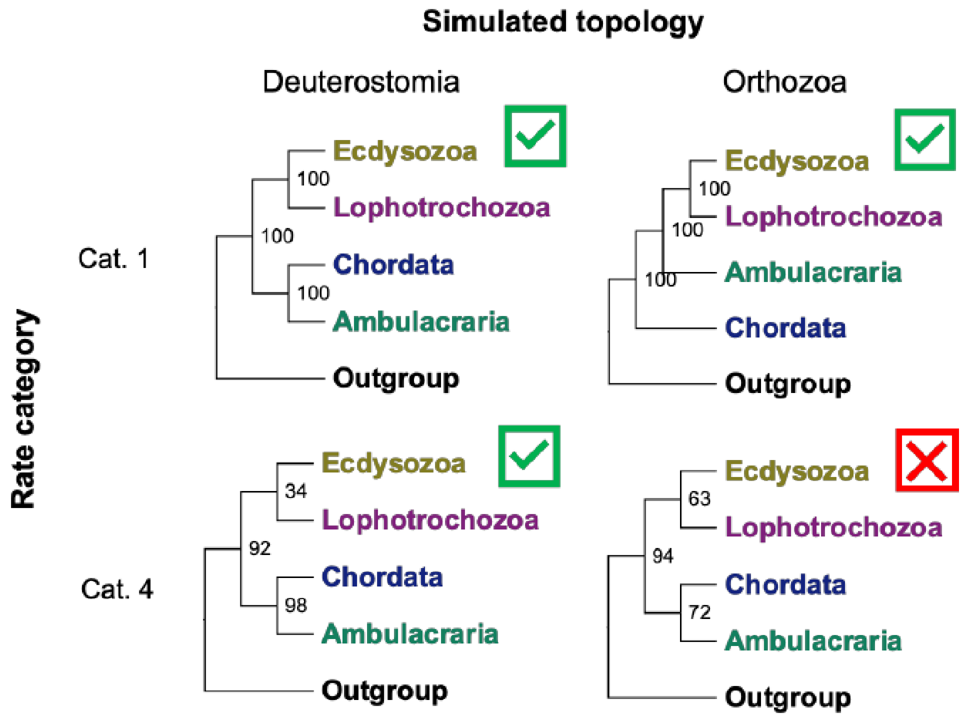


**Fig. S4.**
Likelihood analyses under the LG+F+G4 model of data simulated under an Orthozoa tree and the CAT-Poisson model show that fast-evolving sites (category 4) incorrectly support monophyletic Deuterostomia. The analyses of data simulated with slow-evolving category 1 sites always recovers the 'true' topology.

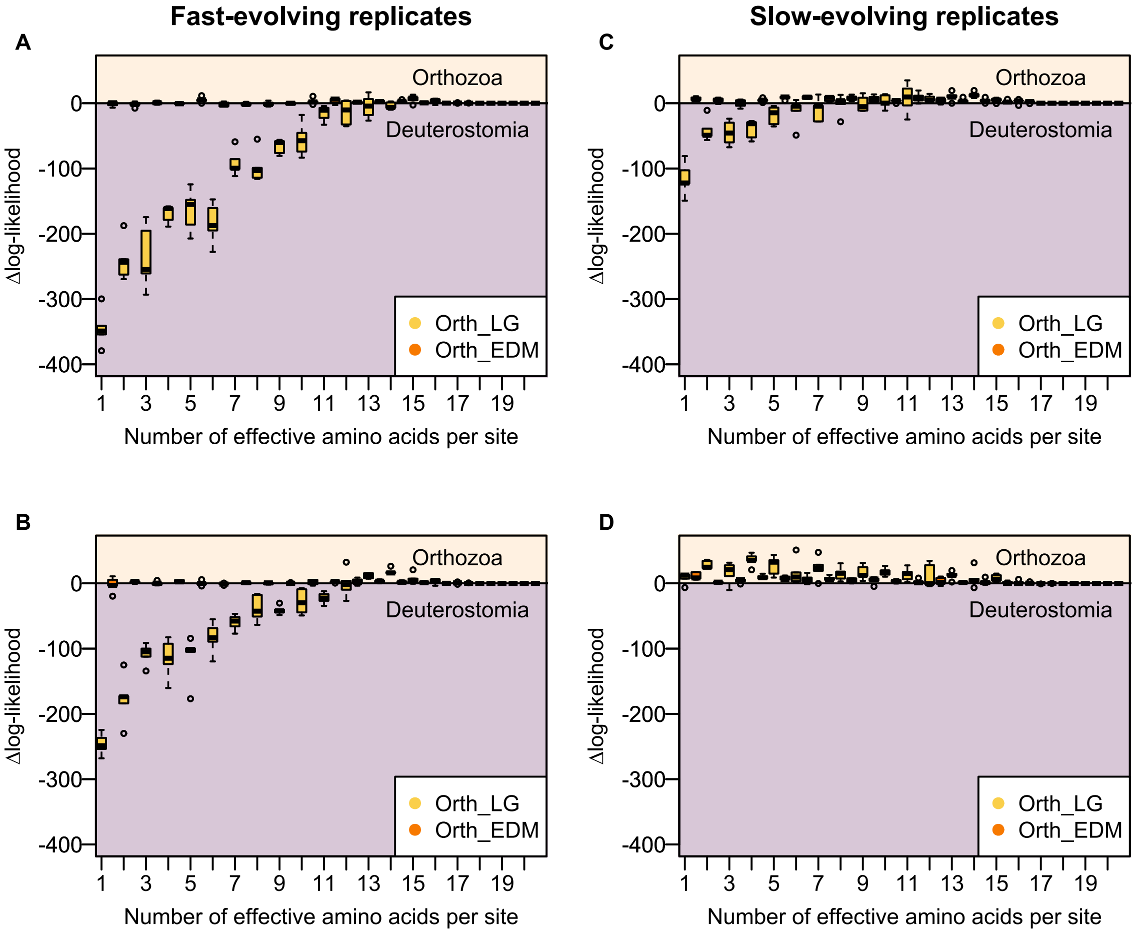


**Fig. S5.**
Simulations under a Deuterostomia topology and the CAT-Poisson model. We show the site-specific Δlog-likelihood between every pair of alternative deuterostome topologies, binned by their effective amino acid number ($k_{eff}$) and under the site-homogeneous LG+G4 (yellow boxes) and site-heterogeneous EDM+G4 (orange boxes) models. For the fast-evolving jackknife replicates, Deuterostomia is supported over Orthozoa, by both models for the jackknife alignment yield the highest (**A**) and lowest (**B**) support for monophyletic Deuterostomia. The site-homogeneous LG+G4 model shows a stronger preference for monophyletic Deuterostomia than the site-heterogeneous EDM model. For the slow-evolving jackknife replicates, Deuterostomia is still supported over Orthozoa under both models and for the jackknife showing the highest support for Deuterostomia (**C**) and for the jackknife supporting Orthozoa (**D**) -- scored under LG+F+G4. For all plots, boxes in the light purple area support Deuterostomia, in the salmon Orthozoa. The black bars correspond to the median Δlog-likelihoods.

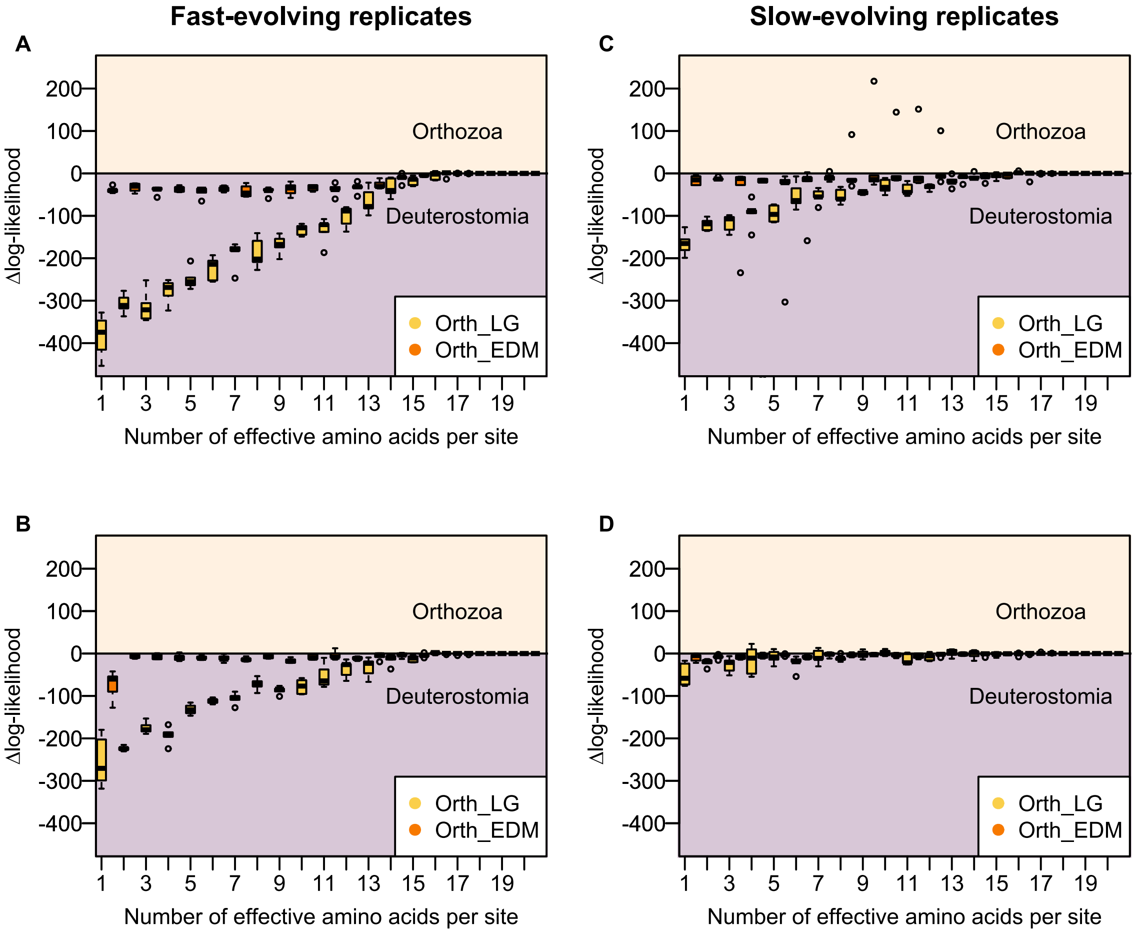


**Fig. S6.**
Simulations under an Orthozoa topology and the CAT-Poisson model. We show the site-specific Δlog-likelihood between every pair of alternative deuterostome topologies, binned by their effective amino acid number ($k_{eff}$) and under the site-homogeneous LG+G4 (yellow boxes) and site-heterogeneous EDM+G4 (orange boxes) models. For the fast-evolving jackknife replicates, Deuterostomia is supported over Orthozoa under the site-homogeneous LG+G4 model, but the site-heterogeneous EDM model always supports the 'true' Orthozoa topology, for both the jackknife alignment with the highest (**A**) and lowest (**B**) support for monophyletic Deuterostomia. The site-homogeneous LG+G4 model shows a stronger preference for monophyletic Deuterostomia than the site-heterogeneous EDM model. For the slow-evolving jackknife replicates, Deuterostomia is still supported over Orthozoa under the LG model for the dataset simulated from an empirical alignment strongly supporting Deuterostomia (**C**). However, LG+G4 supports the 'true' Orthozoa topology for the empirical alignment supporting Deuterostome paraphyly (**D**). The site-heterogeneous EDM+G4 model always supports the 'true' Orthozoa topology. For all plots, boxes in the light purple area support Deuterostomia, in the salmon Orthozoa. The black bars correspond to the median Δlog-likelihoods.

Table S1.
Proportion of alignments and supporting each topology under distinct sampling and site-modelling parameters

| Topology supported by alignments | Measure | Long-branched taxa LG model | Long-branched taxa EDM model | Short-branched taxa LG model | Short-branched taxa EDM model |
|---|---|---|---|---|---|
| Deuterostomia | Topology log-likelihood | 100 | 100 | 99 | 99 |
| | Number of sites | 100 | 99 | 49 | 47 |
| | Average number of sites | 30849 | 23658 | 15650 | 15091 |
| Orthozoa | Topology log-likelihood | 0 | 0 | 1 | 1 |
| | Number of sites | 0 | 0 | 49 | 52 |
| | Average number of sites | 13514 | 12645 | 16643 | 15574 |
| Centroneuralia | Topology log-likelihood | 0 | 0 | 0 | 0 |
| | Number of sites | 0 | 1 | 2 | 1 |
| | Average number of sites | 13350 | 14149 | 12806 | 12809 |

**Data S1. (separate file)**

File dataProvenance.xlsx in DRYAD repository https://doi.org/10.5061/dryad.t76hdr89k
